# Bellymount-Pulsed Tracking: A Novel Approach for Real-Time In vivo Imaging of *Drosophila* Abdominal Tissues

**DOI:** 10.1101/2024.03.31.587498

**Authors:** Shruthi Balachandra, Amanda A. Amodeo

## Abstract

Quantitative live imaging is a valuable tool that offers insights into cellular dynamics. However, many fundamental biological processes are incompatible with current live imaging modalities. *Drosophila* oogenesis is a well-studied system that has provided molecular insights into a range of cellular and developmental processes. The length of the oogenesis coupled with the requirement for inputs from multiple tissues has made long-term culture challenging. Here, we have developed Bellymount-Pulsed Tracking (Bellymount-PT), which allows continuous, non-invasive live imaging of *Drosophila* oogenesis inside the female abdomen for up to 16 hours. Bellymount-PT improves upon the existing Bellymount technique by adding pulsed anesthesia with periods of feeding that support the long-term survival of flies during imaging. Using Bellymount-PT we measure key events of oogenesis including egg chamber growth, yolk uptake, and transfer of specific proteins to the oocyte during nurse cell dumping with high spatiotemporal precision within the abdomen of a live female.

**SUMMARY STATEMENT:** Bellymount-Pulsed Tracking is a novel tool to capture in vivo dynamics for longer duration in *Drosophila* oogenesis.

## INTRODUCTION

Advances in live imaging have been essential for unraveling the dynamics of key molecular events during a variety of biological processes. However, not all processes are equally amenable to live imaging. Capturing very small or rapid molecular events requires sophisticated microscopy (Fuhrmann et al., 2022). Conversely, processes occurring over longer timescales such as growth and development require preserving tissue health during long-term imaging (Martin et al., 2018; Morris and Spradling, 2011; Peters and Berg, 2016; Tsao et al., 2016). *Drosophila* oogenesis is one process that has proven to be resistant to long-term live imaging despite a wealth of genetic tools and molecular characterization (King, 1970; Spradling, 1993; Shimada et al., 2011; Lebo and McCall, 2021; Morris and Lehmann, 1999; Avilés-Pagán et al., 2020; Imran Alsous et al., 2021; Jackson et al., 2023). The major challenge in capturing the journey from a germline stem cell (GSC) to a fertilizable egg is that the process lasts about 7 days and happens inside the female abdomen (He et al., 2011). The GSCs reside in the germarium at the anterior tip of the ovary. Differentiating cells leave the stem cell niche and undergo 4 rounds of incomplete mitosis to generate a 16-cell cyst. One cell is specified as the oocyte and the rest become nurse cells which will endocycle to produce the bulk of the materials required for oocyte growth. The 16 germline cells are encapsulated by a layer of somatic follicle cells to form an egg chamber. The egg chamber exists the germarium and grows several orders of magnitude in volume while undergoing a stereotyped procession of developmental milestones over a period of about 3 days (Dapples and King, 1970; King, 1970; He et al., 2011; Diegmiller et al., 2021). The developing egg chambers migrate posteriorly down the length of the ovary following one another in an assembly line called an ovariole. Each ovariole contains 5-7 egg chambers at any given time and a pair of ovaries will have 32-45 ovarioles (David and Merle, 1968; King, 1970).

Historically, most studies of *Drosophila* oogenesis have been performed using fixed samples which necessarily limit the ability to resolve dynamic processes (King, 1970; Spradling, 1993; Diegmiller et al., 2021; Imran Alsous et al., 2021; Weichselberger et al., 2022). However, as early as the 1970s efforts have been made to develop ex-vivo culture protocols for late-stage egg chambers (Petri et al., 1979; Gutzeit and Koppa, 1982). Optimization of late-stage cultures allows survival of stage 10-14 egg chambers for up to 10 hours (Peters and Berg, 2016; Parsons et al., 2023; Zhang and Liu, 2024). Stage 9 and earlier egg chambers have proven to be more difficult to culture and require the addition of insulin to the media to achieve survival for 4-5 hours (Prasad and Montell, 2007). Further refinement of the culture media allowed for the survival of the germarium to a maximum of 13 hours with follicle cell division rates declining after 6-8 hours (Morris and Spradling, 2011). Nevertheless, the specific growth rates in these culture conditions have not been reported and to date, many live-imaging studies are limited to only a few hours before egg chambers stall their growth and begin dying (Inaki et al., 2022; Zajac et al., 2023). Collectively, these techniques allow a short window to observe processes including the germline-soma coordination in the germarium (Morris and Spradling, 2011; Wilcockson and Ashe, 2021), shuttling of biomolecules across the egg chamber (Shimada et al., 2011; Lu and Gelfand, 2022), follicle cell morphology and migration (Prasad and Montell, 2007; Haigo and Bilder, 2011; Lei et al., 2023; Williams et al., 2022; Miao et al., 2022; Jackson et al., 2023a), and nurse cell changes during later stages of oogenesis (Yalonetskaya et al., 2020; Imran Alsous et al., 2021; Jackson et al., 2023a). Recently, Marchetti et al developed a revolutionary multi-organ co-culture system that successfully maintains egg chamber integrity for up to 3-4 days. However, growth rates remain substantially lower than in vivo and egg chambers fail to progress through developmental landmarks at the appropriate pace (Marchetti et al., 2022).

The recently developed Bellymount tool by Koyama et al (2020) allows visualization of the internal structures including the midgut, crop, intestinal bacteria and ovaries without the need to open the fly’s abdomen by affixing the fly between two glass coverslips using transparent, nontoxic Elmer’s glue. In the original protocol flies were imaged over several days and were released from the mount in between image acquisitions. Though this technique captures unperturbed cellular dynamics in vivo with unprecedented resolution, the need to repeatedly capture and re-mount flies prevents tracking in tissues that lack morphological guideposts for image alignment. This is particularly problematic for oogenesis since each female has dozens of egg chambers, each growing rapidly and moving posteriorly during developments. To overcome this challenge, we developed Bellymount-Pulsed Tracking (Bellymount-PT) by combining the original Bellymount restraint with pulsed anesthesia, and a liquid diet to allow long-term imaging and tracking of abdominal structures for periods up to 12-16 hours. The prolonged imaging duration enabled us to track egg chamber growth, follicle cell movements and visualize transitions between developmental stages. Further, we were able to quantify aspects of yolk uptake and histone accumulation during nurse cell dumping, which have never been previously observed in an in vivo system. This study provides a novel technique for generating insight into the key events of oogenesis in an in vivo system with minimal experimental perturbation.

## RESULTS AND DISCUSSION

### Bellymount-Pulsed Tracking integrates image acquisition with anesthesia to allow long-term imaging in the mount

To maintain long-term tissue health in the endogenous physiological context during imaging we developed Bellymount-PT. Flies were secured between glass surface of a MatTek dish and a compression coverslip using Elmer’s glue and 0.48 mm adhesive spacers (Fig. 1A). Flies were repeatedly anesthetized using CO_2_ to prevent voluntary and involuntary movements during image acquisition, CO_2_ pulsing allowed the flies to awaken and feed between timepoints (Poinapen et al., 2017; Koyama et al., 2020; Tang et al., 2021). Since CO_2_ anesthesia adversely affects fertility in flies (Shen et al., 2020) we minimized the duration of anesthesia by integrating image acquisition with CO_2_ pulsing via an Arduino controller. The Arduino opened a solenoid valve on the CO_2_ tank in response to a signal from the image acquisition software (Fig. 1B) 2 minutes before the beginning of imaging and closed the valve immediately after image acquisition. Pulsed anesthesia allowed flies to awaken, feed, defecate, and lay eggs while restrained in the mount between image acquisitions (Fig. 1A,C,D). We tracked egg chambers with imaging intervals from 10 min to 2 hours depending on the experimental requirements.

**Fig. 1.**
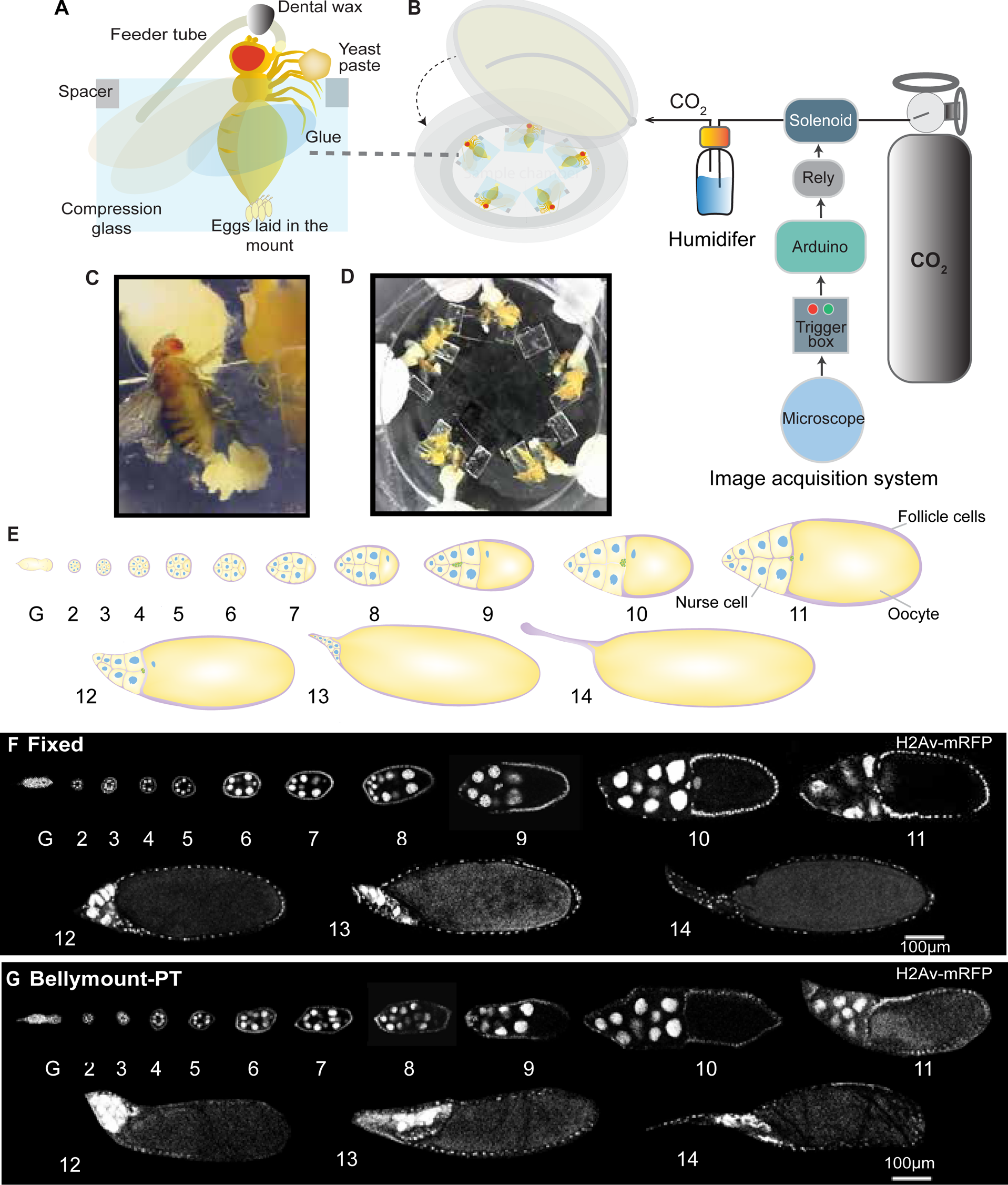
Bellymount-Pulsed Tracking (PT) allows long-term live imaging of oogenesis in live females. (A) Schematic of a fly mounted for Bellymount-PT. Mated and well-fed females were glued on the glass surface of a MatTek dish using Elmer’s glue and compressed between the dish and the glass coverslip with 0.48 mm spacers to secure the fly during imaging. A small amount of fresh yeast paste and a feeder tube containing custom liquid apple juice, yeast extract food ensure proper nutrition. (B) Image acquisition setup for confocal microscope and pulsed CO_2_ anesthetic. An Arduino programmed to regulate the CO_2_ flow via a solenoid value is connected to a signal from the microscope. CO_2_ is pulsed into the sample dish for four minutes during acquisition through a humidifying chamber. Flies are allowed to awaken and feed between time points. (C) A female that remained alive and fecund after 16 hours in the mount without imaging. Note the large number of eggs which have accumulated to the posterior of the fly. (D) Whole mount with flies and feeder tubes. Five females can be imaged in the same dish simultaneously. (E) Schematic showing oogenesis stages. The blue circles represent the nurse cells and the oocyte nuclei, the pink border represents the follicle cell layer, and the cells in green represent border cells. (F) Staged fixed egg chambers expressing H2Av-mRFP from multiple females imaged at 20x (0.8 NA) on Zeiss LSM980. (G) Collection of staged live egg chambers expressing H2Av-mRFP from multiple females acquired using Bellymount-PT imaging on the same microscope as F. These egg chambers were imaged through the intact cuticle of the female abdomen. Adjacent egg chambers were cropped out and images were processed for presentation using background subtraction, gaussian filter and median filters in Fiji.

Up to 5 flies can be mounted on a single 50 mm dish (Fig. 1B,D; Movie 2). We typically imaged 3-4 flies during each session and the unimaged flies served as controls to determine the effects of laser exposure on fly health (Fig.2A,B). This ability to image multiple flies in a single experiment will help to control for day-to-day variability when multiple genotypes are imaged simultaneously. Since oogenesis is extremely sensitive to the nutritional status of the female (Terashima and Bownes, 2004; Shimada et al., 2011), we developed a custom apple juice and yeast extract liquid food which was provided via a cotton wick in a bent capillary feeder tube (Fig.1A, S1). To stimulate olfactory feeding cues, we included fresh yeast paste and banana baby food within the reach of each fly’s legs. The food and a moist absorbent pad positioned at the top of the imaging chamber kept the flies hydrated during imaging (Fig. 1A-D).

**Fig. 2.**
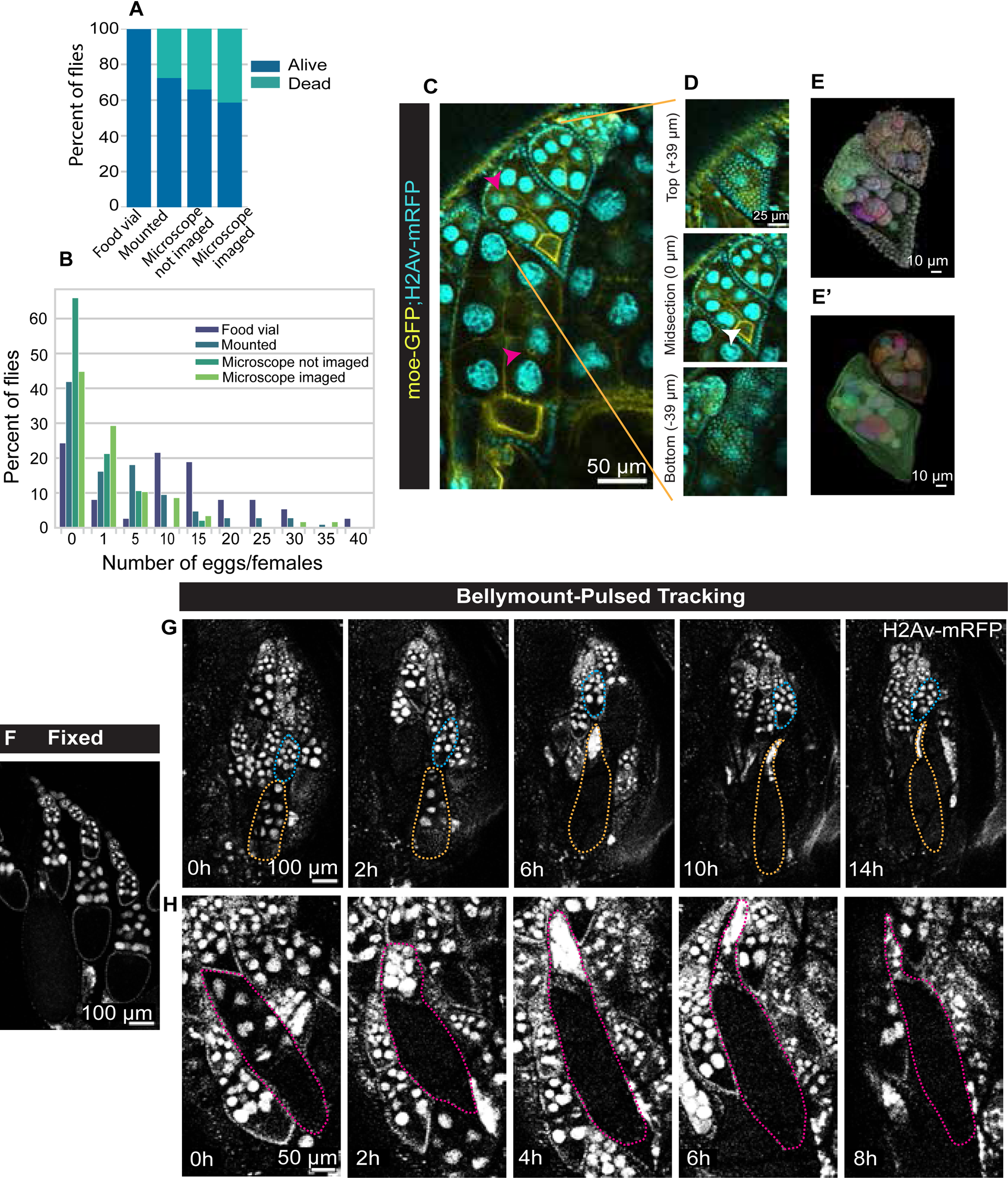
Pulsed anesthesia allows tracking for over 14 hours. (A) Survival percentage for females after 16 hours of imaging compared to various controls. To understand the effect of mounting, anesthesia, and laser exposure we compared imaged flies (n=58) to controls that were either never placed in the Bellymount-PT chamber (food vial, n=37); mounted, but never exposed to the imaging protocol (mounted, n=105), and those that were placed on the microscope during imaging, anesthetized but not exposed to laser (microscope, not imaged n=47). Most of the observed mortality can be attributed to the restraints, though anesthesia and exposure to laser further decrease survival. (B) Histogram of eggs laid in 16-hour period for the same flies as in A. Females laid 0-40 eggs during the imaging period. Flies that remained in their food vials laid an average of 10 eggs per female while flies imaged laid 2 eggs on average. (C) Single time point (not time-lapse) representative image of multiple egg chambers within a single ovariole expressing H2Av-mRFP and Moesin-GFP using Bellymount-PT acquired on Olympus multiphoton microscope (25x water objective). (D) All three cell types (oocyte, nurse, and follicle cells) can be visualized through the cuticle. There is sufficient z-resolution to ∼ 80 µm to capture the full volume of egg chambers stage 8 and younger. Pink arrowheads indicate ring canals and white indicates oocyte nucleus. Insets represent the top (+39µm), midsection (0µm), and bottom (-39µm) z-slices of a stage 6 egg chamber. (E,E’) 3D reconstruction of the egg chambers in C with nuclear H2Av-mRFP signal and the 3D constructed surface. (F) Fixed egg chambers with the same genotype and imaging as G-H for comparison. (G-H) Live, developing egg chambers expressing H2Av-mRFP imaged over time using Bellymount-PT. (G) Multiple egg chambers can be tracked in a single experiment. The egg chamber marked in blue starts in early stage 8 and increases in size over the imaging duration. The egg chamber marked in yellow starts in stage 10 and progresses through stage 14 with complete dorsal appendages formation visible by 14 hours. (H) Egg chamber marked in magenta shows the progression the of egg chamber from stages 10-14. Images in F-D were imaged on Zeiss LSM980 microscope with a 10x air objective. Images in the (C-H) were processed for presentation using background subtraction, gaussian filter and median filters in Fiji.

### Females mounted for Bellymount-PT remain alive and fecund

Limiting the exposure to anesthesia to once every 2 hours and providing continuous access to food allowed ∼60% flies to survive for greater than 16 hours of imaging (Fig. 2A). To understand the causes of mortality during imaging we maintained controls on standard food in vials, in the mount without exposure to CO_2_ or laser, and on the microscope exposed to repeated anesthesia but not imaged. As expected, nearly all flies maintained in vials survived and flies restrained in the mount and exposed to both laser and CO_2_ had the highest mortality. The bulk of the mortality seems to be due to the restraint system itself since the flies in the mount that were not exposed to anesthesia or the laser had only about a 10% increase in survival compared to those that were imaged. We observed that much of the mortality resulted from drowning or dehydration when the wick for the liquid food was not ideally placed above the fly.

Fecundity is a crude reflection of the rate of egg chamber development. To understand the influence of experimental conditions on egg production, we counted the eggs laid by females imaged during an overnight (16 hour) session, along with other controls as described above. Restrained flies frequently continued to lay eggs which accumulated in the mount at the posterior of the fly (Fig. 1C, Movie1-3). On average, flies that remained in their food vials at 19-20℃ laid 10±2 (SEM) (n=37), while flies that were imaged laid 3±1 (SEM) eggs per female overnight (n=59) (Fig.2B). Although restraint did significantly reduce egg production (*p*<0.0001), the fact that egg production continues even during imaging indicates that oogenesis is not completely stalled under the Bellymount-PT protocol.

### Imaging through the cuticle allows single-cell resolution of developing egg chambers

Bellymount-PT allows the identification of all stages of oogenesis from the germarium through stage 14 using H2Av-mRFP, although not all stages were visible in the same female (Fig. 1E-G). The precise imaging parameters required depend on the resolution and duration of imaging required for a specific experiment. We can easily resolve individual H2Av-mRFP nuclei and Moesin-GFP at the cortex within egg chambers to a Z-depth of 75-80 µm using an Olympus multiphoton microscope with a 25x water objective (NA 1.05) (Fig.2C). The nuclear signal of follicle cells defined the outer egg chamber boundary allowing for accurate three-dimensional reconstruction of full egg chamber volumes up to stage 8 (stage 6 shown in the Fig. 2D). Later-stage egg chambers could not be imaged to their full depth, but in all cases, the midsection area functions as a proxy for egg chamber size. We can resolve subcellular structures including ring canals and oocyte chromatin (Fig. 2C-E’; Movie 4-6). At lower magnification (Zeiss 980 with a 10x NA0.5 air objective) several egg chambers of different ages can be imaged simultaneously, (Fig. 2G) increasing the throughput of the technique. The wider range of view of the lower magnification objective also aided in long-term tracking since egg chambers move within the female abdomen throughout development and the female herself moves a small amount within the restraints during imaging intervals. Under these conditions, we can track multiple egg chambers through different stages of development for 8-16 hours (stage 10-14, Fig.2G,H, Movie 7,8; stage 7-8, Fig.3F,H Movie 9,10). At 20x magnification we can visualize border cells migration (Movie 11) and egg chamber rotation (Movie 12 and 13).

**Fig. 3.**
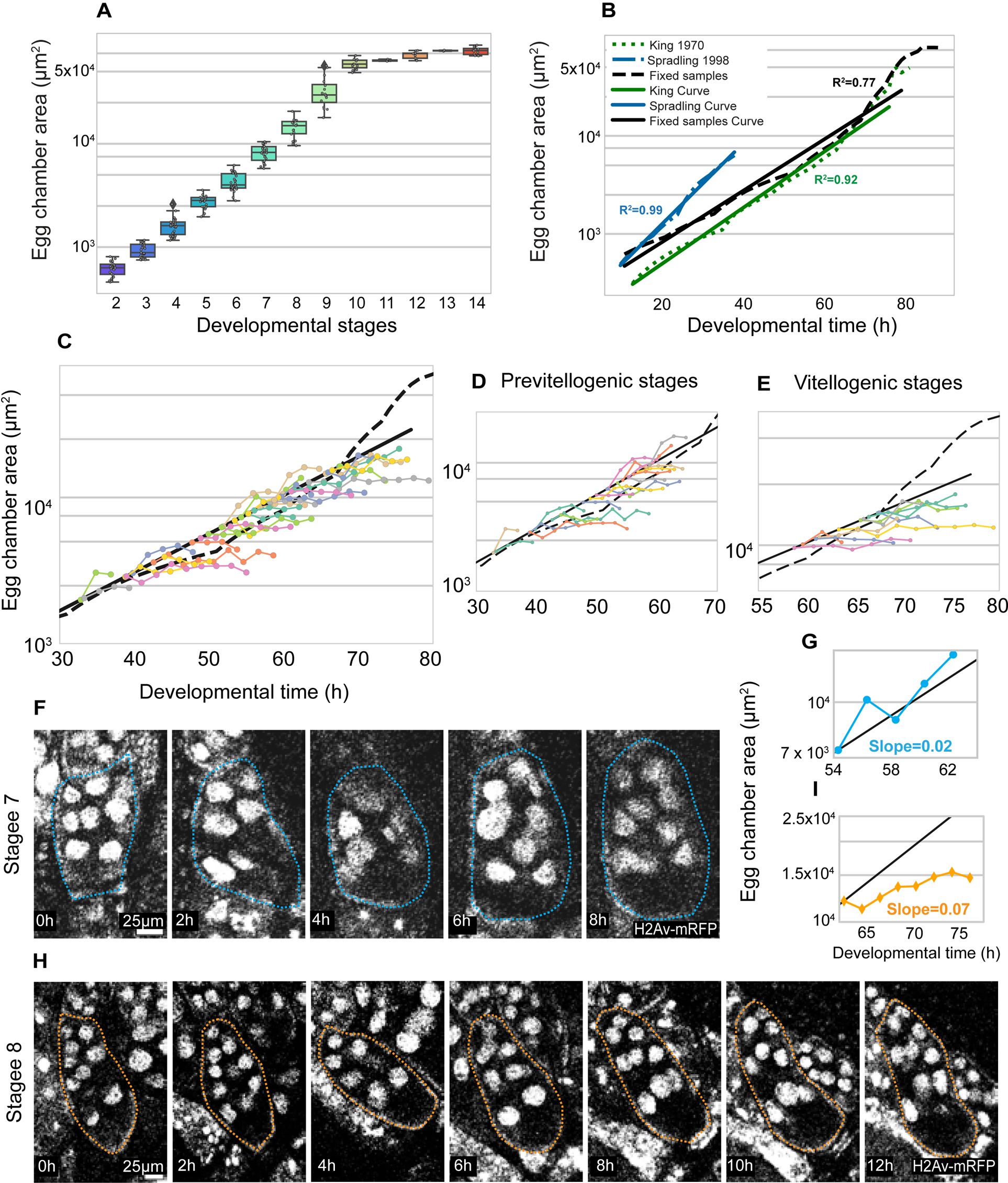
Bellymount-PT allows live monitoring of egg chamber growth. **(A)** The midsection area of fixed egg chambers by stage. Fixed egg chambers were staged based on milestones of oogenesis as per (King 1970; see methods). The measured sizes combined with the frequency of each stage, total number of egg chambers per ovariole, and the number of eggs laid by individual females in 24 hours were used to construct a standard growth curve (black line panel B) for the fly line we used for live growth analysis (w; Mat-ɑ-tub67-gal4/CyO; Mat-ɑ-tub15-gal4, H2Av-mRFP/TM3) (B) Comparison of standard growth curves from this study and previous literature derived from fixed samples. Egg chamber areas from King et al 1970 (green) and Spradling 1993 (blue) were used to generate an exponential standard growth curve along with the corresponding time estimates. (C) The midsection area of 38 egg chambers tracked using Bellymount-PT with 2-hour imaging intervals. The resulting traces were aligned to the standard growth curve shown in B (black) by their initial (t0) area and with subsequent timepoints based on actual image acquisition times. Individual egg chambers were tracked for 4-18 hours. (D) A subset of the area traces in C that begin in the previtellogenic stages (4-7). (E) A subset of the area traces in C that begin at the start of the vitellogenic stages (stage 8). (F) A previtellogenic egg chamber starting at stage 7. (G) The change in midsection area over time for the egg chamber in E compared to the standard growth curve in black. Note the agreement between predicted and observed growth rates. (H) A vitellogenic stage 8 egg chamber tracked for about 12 hours. (I) The growth rate of the egg chamber in H compared to the standard curve. Note that growth is slower than predicted. (F and H) were processed for presentation using background subtraction, gaussian filter and median filters in Fiji.

### Bellymount-PT enables tracking live growth rates of individual egg chambers

Due to the challenges of live imaging oogenesis, existing estimates of egg chamber growth rates are based on the number of egg chambers of each stage in individual females at a given time and the average number of eggs laid per female per day. Moreover, literature estimates differ greatly in the time assigned to each stage due to differences in genetic backgrounds and fly husbandry (David and Merle, 1968; King, 1970; Spradling, 1993). To account for our own strains and fly husbandry, we generated a new growth curve for the developmental time of w; Mat-ɑ-tub67-gal4/CyO; Mat-ɑ-tub15-gal4, H2Av-mRFP/TM3 females maintained at 23±1℃. Note that our standard growth curve is constructed from flies that were kept at warmer temperatures than our imaging conditions. Since the conventional staging of egg chambers is largely based on size and we wanted to measure growth rates, we staged the egg chambers based on size-independent parameters whenever possible and measured the egg chamber midsection area for each stage from the fixed samples (n=15 females; n=234 egg chambers) (Fig. 3A; see methods). We recorded the fecundity (eggs laid/female/day=52±2.7 SEM), egg production rate (1.45 eggs/per ovariole/day), and the frequency of each stage in an ovariole to estimate the duration of each stages 2-14 (Table S1). This data allowed us to estimate the expected growth rate for egg chambers during unperturbed oogenesis which is well fit by an exponential equation (A=273 e ^0.06t^, where t=time in hours and A=area in µm^2^). Our measured standard growth curve is within the range reported in previous literature (King, 1970; Spradling, 1993) (Fig.3B).

Next, we sought to measure the real-time growth rate of individual egg chambers inside the female abdomen using Bellymount-PT. We were able to track the growth of egg chambers which were initially in stages 4-10. We measured the midsection area of each egg chamber over time. To test if the growth rates observed using Bellymount-PT matched those expected from the fixed growth curve we aligned each trace to the growth curve based on the initial area with the subsequent times based on the imaging intervals. While some egg chambers had growth rates that aligned well with the expected growth curves (Fig. 3C-G, Movie 9), most grew more slowly than our standard (Fig. 3C-E, H,I, Movie 10). During the initial 2 hours of imaging, the imaged egg chambers reached an average of 95% of the expected size, however by 8 hours, they had only reached 72% (Fig. S2). We note that although growth rates decreased, many of the size-independent aspects of oogenesis progressed at a relatively normal rate indicating that the egg chambers continued to develop. This suggests that the midsection area may be an underestimate of egg chamber volume when the egg chambers remain within the female abdomen compared to fix samples compressed between two coverslips. In particular, stage 10 egg chambers were often seen to progress to stage 14 in about 10 hours as expected but did not appear to grow as much as expected (Fig.2 GH).

### Bellymount-PT allows unprecedented visualization of the events of oogenesis

Finally, we employed Bellymount-PT to capture dynamic protein localization events during oogenesis. First, we focused on the uptake of yolk protein into the oocyte from the hemolymph. The initiation of yolk uptake is a checkpoint for further progression of the egg chamber development. Active yolk uptake starts from early stage 8 and continues through stage 10 (Terashima and Bownes, 2004; Shimada et al., 2011; Row et al., 2021; Isasti-Sanchez et al., 2021). Inadequate nutrition and environmental stress stall oogenesis at stages 7 or early 8 by preventing yolk uptake (Terashima and Bownes, 2004; Shimada et al., 2011; Hara and Yamamoto, 2022). *Drosophila* yolk proteins (Yp1-3) are predominantly made in fat bodies and ovarian follicle cells, and their endocytosis by the oocyte is stimulated by factors including hormonal levels (Richard et al., 1998) and male-derived sex peptide received during mating (Soller et al., 2006). Thus, yolk accumulation requires the coordination of multiple organ systems and external factors. This explains the difficulty in maintaining egg chamber growth during these active vitellogenic stages in ex vivo culture systems (Peters and Berg, 2016). To test if Bellymount-PT supports yolk uptake, we tracked Yp1:sfGFP (Hara and Yamamoto, 2022) accumulation in a stage 8-10 egg chambers (n=6). Since in later stages growth is dependent on yolk uptake, we aligned the growth curves to the expected initial age based on the initial oocyte percentage instead of the midsection area (Fig. 4C). We used an endogenously tagged H2Av-mCherry to manually segment the oocyte and measured the total amount of Yp1-sfGFP protein in ½ of the egg chambers over time. We found that yolk continually accumulates at a relatively constant rate for the duration of the measured stages (Fig. 4A,B, Movie 14). These measurements of yolk uptake from the hemolymph are only possible by preserving the physiological context inside the intact female abdomen using Bellymount-PT.

**Fig. 4.**
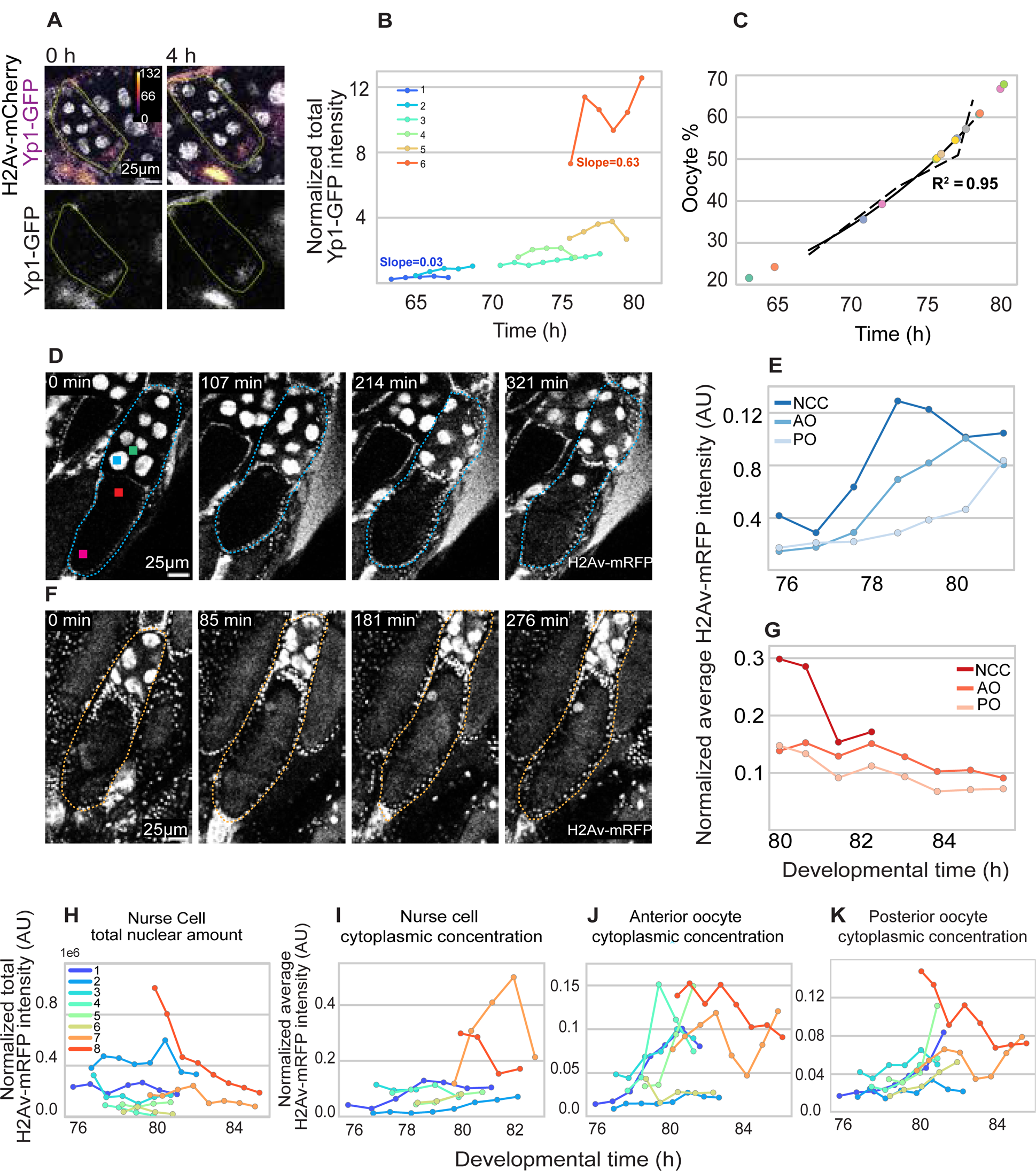
Bellymount-PT allows for the observation of dynamic events during oogenesis. (A) Yolk protein accumulation during vitellogenic stages was tracked in flies expressing Yp1:sfGFP. (B) Quantification of the total Yp1:sfGFP intensity in the oocyte over time for 6 different egg chambers. We observe a steady increase in Yp1:sfGFP in all oocytes (Egg chamber ages were estimated as in (C)). (C) A growth curve was generated based on the oocyte percent to estimate the age of the egg chambers to understand the timeline of yolk uptake and histone accumulation (see methods and Fig. S3A for correlation between egg chamber area and oocyte percent). (D) Stage 10 egg chamber expressing H2Av-mRFP during early nurse cell dumping. Note the flow of H2Av from the nurse cells into the oocyte. Squares indicate the region of the egg chamber quantified in E,G-K. (E) Quantification of average pixel intensities in the indicated regions of D normalized to the intensity of the follicle cell nuclei (NCC-Nurse cell cytoplasm; AO-Anterior oocyte cytoplasm; PO-posterior oocyte cytoplasm). Note the steady increase of H2Av in all three regions as it exits the nurse cell nuclei (F) A more advanced, stage 11 egg chamber during later stages of dumping. (G) Quantification of the H2Av intensity in different regions of the egg chamber shown in F over time. Note that H2Av declines in all regions at these later stages. (H-K) Changes to the normalized H2Av intensities in various parts of 8 independent egg chambers of various ages. Each color corresponds to the same egg chamber in all panels. Egg chamber ages were estimated from the standard growth curve in C based on the percent of egg chamber area occupied by the oocyte at t0. Together these data are consistent with H2Av being lost from nurse cell nuclei and flowing gradually posteriorly into the oocyte. (H) The total integrated nurse nuclear intensities represent the total amount of H2Av in the nurse cell nuclei over time. These intensities show a consistent decline over the course of nurse cell dumping indicating that H2Av is being lost from nurse cell nuclei. (I)The mean cytoplasmic intensity of the nurse cells indicates an initial increase followed by a decline in H2Av nurse cell cytoplasmic concentration. (J), The intensity in the anterior oocyte shows an initial dramatic increase followed by a slight decline (K), The intensity in the posterior oocyte is initially flat and increases only later during dumping. The accumulation is delayed compared to the anterior oocyte in J indicating an overall posterior flow. Images in (A D, F) were processed for presentation using background subtraction, gaussian filter and median filters in Fiji.

We were interested in measuring maternal protein transfer from the neighboring nurse cells to the oocyte during dumping. Histones are essential chromatin components that are preloaded into the oocyte during oogenesis to support the early embryonic cell cycles (Woodland and Adamson, 1977; Günesdogan et al., 2010; Stephenson et al., 2021; Shindo et al., 2022). The oocyte is unusual in that it generates large stores of both replication-dependent and replication-independent histones, which are stable beyond S-phase (Li et al., 2012). Maternal histone loading is of great importance, as histones concentration regulates the progression of early embryonic development (Amodeo et al., 2015; Shindo and Amodeo, 2019; Chari et al., 2019; Shindo and Amodeo, 2021). Like most proteins, replication-dependent histones are synthesized in the nurse cells and transferred to the oocyte during the later stages of oogenesis (Ambrosio and Schedl, 1985; Ruddell and Jacobs-Lorena, 1985; Walker and Bownes, 1998). However, a recent study showed that H2Av is transferred to the oocyte as early as stage 9 (Stephenson et al., 2021), suggesting that different histones can behave differently during oogenesis.

To measure the flow of histones into the oocyte we measured the concentration of H2Av-mRFP in various regions of the egg chamber during nurse cell dumping (Fig. 4D-K). We also segmented the nurse cell nuclei to measure how the nuclear volume and total H2Av-mRFP changed over time (Fig. S3). We found that the nurse cell nuclei lost H2Av-mRFP over time (Fig 4H). However, nuclear volume decreased more rapidly than the H2Av-mRFP loss resulting in a transient increased nuclear concentration (Fig. S3). At the same time, the cytoplasmic intensities of H2Av-mRFP increased in the nurse cells and the anterior and posterior regions of the oocyte with the anterior preceding the posterior (Fig. 4D-E, Movie 15). After completion of dumping, we observed a drop in H2Av-mRFP concentrations both in the anterior and posterior regions of the oocyte (Fig. 4FG, Movie 16), likely due to the dynamic streaming of the oocyte cytoplasm and an increase in the overall volume of the oocyte. Bellymount-PT enabled us to capture the dynamic flux of the maternal proteins in real-time during the later stages of oogenesis

Along with the above key milestones of oogenesis we captured instances of border cell migration (Movie 11), and rotation of egg chamber of in wide range of egg chamber stages (Movie 12,13). Though egg chamber rotation is well documented in stage 5-8 egg chambers (Haigo and Bilder, 2011; Horne-Badovinac, 2014) it appears to continue into stage 9-10 when egg chambers are maintained in vivo (Movie 12, 13). We also observed a novel behavior in egg chambers during stage 12-13 where they appear to slip sidewise rather than a complete rotation (Movie 13). The ability to observe these multiple aspects of egg chamber development highlight the usefulness of Bellymount-PT to explore oogenesis in physiological context.

### Conclusion

Bellymount-PT provides a powerful means to capture dynamic processes in living animals with minimal perturbation. We have focused on *Drosophila* oogenesis, which is intricately regulated by organismal signals including nutritional availability, mating status, and environmental conditions. While egg chambers can survive in ex vivo culture for a limited time, these systems are unable to faithfully replicate the physiological growth cues, signaling events, and stress response systems found in vivo. The non-invasive nature of Bellymount-PT allows access to internal, sub-cellular details while preserving the native physiological milieu. We can capture all stages of oogenesis and track multiple egg chambers for over 16 hours, observing progressive developmental stages. The ability to image vitellogenesis, a critical checkpoint in oogenesis that is difficult to study in cultured systems, demonstrates the improved egg chamber health compared to other state of the art techniques.

One notable constraint of Bellymount-PT is the inability to capture the entire length of oogenesis with normal growth rates. We observed a ∼40% mortality rate and decreased fecundity compared to unrestrained flies. A major cause of mortality was drowning in the liquid food within the mounts. Further optimization of the feeding protocol may improve fly health and thereby increase the overall rate of egg chamber development. Another hurdle for fly health during imaging is presented by the common practice of housing microscopes in cool, dark rooms. Increasing the temperature and scheduling imaging sessions to align with the fly circadian rhythm would also likely further increase fecundity. Nonetheless, Bellymount-PT represents a significant improvement in the length of egg chamber growth and survival over existing techniques. Bellymount-PT can be easily implemented in any laboratory equipped with a confocal microscope without requiring specialized equipment. The capability to image multiple females simultaneously increases the throughput and allows multiple genotypes to be imaged under identical conditions.

The utility of Bellymount-PT extends beyond oogenesis. Indeed, the original Bellymount protocol was developed to capture intestinal stem cell dynamics. Within the gut Bellymount has sufficient resolution to differentiate individual bacteria (Koyama et al., 2020). Bellymount-PT is compatible with the long-term imaging of other abdominal tissues such as the crop, tracheal tubes, fat bodies, and blood cells. Therefore, Bellymount-PT presents an attractive alternative to current ex vivo culture modalities, especially for processes such as yolk uptake that require coordination across multiple tissues.

## MATERIAL AND METHODS

### Fly stocks

*Drosophila* stocks were maintained at room temperature (22±2°C) on standard molasses corn fly media. The following fly stocks were used in this study: w; Mat-ɑ-tub67-gal4, Moesin-GFP/CyO; Mat-ɑ-tub15-gal4, H2Av-mRFP/TM3 (BDSC, 31775) (recombined line received as gift from He Lab, Dartmouth College, NH, USA), w; Mat-ɑ-tub67-gal4/CyO; Mat-ɑ-tub15-gal4, H2Av-mRFP/TM3 (Wieschaus Lab, Princeton University, USA) (Hunter and Wieschaus, 2000), yw; Sp/CyO; EndoH2Av-mCherry/TM3 (a generous gift from Yuki Shindo) (Shindo and Amodeo, under preparation), Yp1-sfGFP/CyO (VDRC, 318746) (Hara and Yamamoto, 2022).

### Fly mount preparation

The Bellymount imaging system was adapted from Koyama et.al. (2020). 4-5day old females raised on standard molasses corn fly food supplemented with fresh yeast paste were transferred to apple juice agar plates with fresh yeast paste 12-20 hours before imaging. Healthy and well-fed females were anesthetized on a fly CO_2_ pad for mounting. A maximum of five females per experiment were mounted on the glass surface of a 50 mm MatTek dish (MatTeck, P50G-1.5-30-F) using clear Elmer’s Liquid School Glue (Elmer’s, E309). Adherence of the flies to the dish was accomplished as described in Koyama et.al. (2020). A small drop of Elmer’s glue was placed on the MatTeck dish near the edge of the glass surface. An anesthetized female was carefully placed on the glue and pressed gently with blunt forceps. The fly was secured by placing a 0.5 cm^2^ compression glass starting from her thorax (cut from a 24 x 60 mm cover glass; VWR International, 48393-106). A pair of 2 mm^2^ double-sided adhesive spacers of 0.48 mm thickness (Millipore-Sigma, GBL620004-1EA) were placed on either side of the fly between the compression glass and the dish surface (Fig.1A).

Flies were provided with a novel liquid food to keep them hydrated and well-fed during long-term imaging. A feeder tube was constructed by bending a 2 cm long glass disposable micropipette (VWR International, 53432-921) into a “J” shape using a Bunsen burner. This shape allowed a larger volume of liquid food to be provided to each fly within the space confines of the imaging dish. A wick made from cotton (Equate beauty, 681131169684) was inserted into the short end of the J to allow for the slow release of liquid media. Feeder tubes were filled with freshly made 0.5 mg/ml yeast extract (Thermo Fisher Scientific, BD 212750) in undiluted apple juice (Ocean Spray, 10031200007206). A few grains of active yeast (MP, 02101400-CF) were added liquid food, and the tip of the wick was dipped in banana baby food (Gerber, 015000076054) immediately before mounting. The feeder tubes were secured with dental wax (Fresh Knight, 40201616938) adjacent to each fly’s proboscis so they could feed during imaging intervals. Care was taken to position the feeder tube at an appropriate distance to avoid drowning the fly. A small dollop of yeast paste was provided near the flies’ legs to stimulate feeding.

### Pulsed Anesthesia

For effective anesthetization and to maintain optimum humidity essential for long-term imaging, we covered the sample dish with a 50 mm petri dish lid (Fisher Scientific, 351007). We used this lid instead of the lid of the MatTek dish to increase the depth of the sample dish to allow insertion of moistened absorbent pads (Hazmat Sorbent Pads, S-14748) cut to a diameter of 40 mm with a slit of 15 mm along the diameter and CO_2_ tubing. Both were secured with a metal mesh (Genesee Scientific, Flystuff 57-100 Mesh, Stainless Steel 97um) cut to 50 mm diameter. A hole was drilled on the side wall of the lid to allow entry of a 1/16” diameter flexible plastic tube (Millipore Sigma, Z765228) for the administration of CO_2_ anesthesia. Through this hole, CO_2_ tubing was inserted and glued using KWIK-SIL adhesive silicone glue (World Precision Instruments, 60002) to the wall of the dish. 100% CO_2_ at 5 PSI (Airgas®, AS215A320) was bubbled through distilled water in a 500-mL PeCon humidification bottle (PeCon) to increase humidity when CO_2_ was administered.

An Arduino (Arduino, 7630049200623) was programmed to open a solenoid valve (Plum Garden, PL-220101) via a rely (TOLAKO, BJ-DT0Y) on receiving a trigger signal from the microscope for 4 minutes. All time-lapse imaging experiments were conducted on an inverted Zeiss LSM980 microscope controlled by Zen blue (Carl Zeiss, Zen version 3.6). Zen’s Experiment Designer module was used to coordinate trigger out signals via an external trigger box (Carl Zeiss, User Interface Box 1437-440) to the Arduino. This module also allowed us to introduce imaging pause and acquisition intervals. At the start of the experiment, we set a trigger out signal to Arduino. Upon receiving the signal from the microscope the Arduino opened the solenoid valve for 4 minutes allowing CO_2_ flow into the sample dish. After sending out the trigger-out signal we set a 2 min imaging pause following CO2 inflow. This pause helps the flies succumb to anesthesia before image acquisition. We limited the duration of image acquisition to 2 min to correspond to the anesthesia time. Following completion of imaging the CO_2_ was withdrawn. Imaging intervals were set according to the experimental requirement (2 hours for growth rate measurements (Fig. 2GH, Fig. 3FH), 1 hour for yolk protein uptake (Fig.4A) and 10-11 minutes for the H2Av-mRFP transfer experiment (Fig. 4DF)).

### Microscopy

All the timelapse imaging was conducted on Zeiss LSM980 confocal microscope with Airyscan. The system is equipped with four lasers (405nm, 488nm, 561nm, and 633nm), 2-multi-alkali PMT detectors, a GaAsP detector, and a motorized Axio Observer-7 stage with a z-piezo insert. Images were acquired in Zeiss Multiplex mode (Airyscan MPLX CO-8Y) with either 10x (0.5 NA) air objective or a 20x (0.8 NA) air objective. We were able to reach a Z-depth of ∼100 µm with a Z step size ranging from 3-5µm. We utilized the fastest mode of acquisition with pixel time range 0.34 µs - 0.56 µs for 1-2, fluorophore with a pixel sizes ranging from 0.274 x 0.274 x 5 µm - 0.149 x 0.149 x 4 µm respectively. Further details of pixel size, pixel time, fluorophores used, laser setting, imaging intervals are described for each experiment in Table S2.

Egg chambers used for 3-dimensional reconstructions (Fig. 2CD) were acquired using an upright Olympus FVMPERS multiphoton microscope equipped with the InSight DeepSee Laser System, multi-alkali photomultipliers, two GaAsP detectors, Olympus 25x water immersion objective (XLPLN25×WMP2, NA 1.05 531) running FluoView software. We used 920 nm and 1040 nm lasers for GFP and mRFP respectively. We imaged to a Z-depth of 110µm with 3µm step size. Note, this microscope was not equipped for pulsed anesthesia and used to acquire only single time frame images (Fig. 2C-E’).

Brightfield images and recordings of whole flies in the mounts (Fig. 1C,D, Movie. 1-3) were captured using a digital USB-pluggable 60x-250x Digital microscope (Plugable, USB2-MICRO-250X).

### Fixed egg chamber sample preparation

Ovaries from 5-7 females were dissected in 1% PBS-Tween and fixed in a 1:1 mixture of 4% formaldehyde (Thermo Scientific, 28908) and heptane (Sigma Aldrich, 34873) for 20 min at room temperature. After fixation samples washed 3 times for 10 min each in 0.3% PBS-Tween. Samples were then mounted in a 1:1 mixture of Aqua-Poly/Mount (Polysciences, 18606) and RapiClear (SUNJin Lab, RC147001) mounting media. The experiment was performed twice for a total n=15 females. These fixed samples were used to construct a standard growth curve.

### Image quantification

#### Egg chamber midsection area measurements

Nuclear H2Av-mRFP signal from follicle cells was used to trace the midsection area of the egg chambers using the polygon tool in FIJI (Version 2.14.0/1.54f) (Jia et al., 2016).

#### Staging fixed egg chambers

Where possible we used size independent criteria to stage egg chambers. Due to lack of suitable markers, egg chambers younger than stage 4 were distinguished based on their size. The chromosome condensation status, which changes from blob-like to dispersed, was used to distinguish stage 4, 5, and 6. Oocyte nuclear position, which moves anteriorly, distinguished stage 6 from 7. Increased proportion of the oocyte area distinguished stages 8-11 along with border cell migration and nurse cell dumping. By the completion of stage 8, the oocyte accounts for 30% of the egg chamber area. The initiation of the border cell migration is an indication of stage 9 and completion of migration identified stage 10. The onset of dumping at stage 11 brings about a dramatic increase in the oocyte size, whereas stages 12 and 13 were distinguished based on the appearance of dorsal appendages. Stage 14 is marked by the complete clearance of the nurse cell nuclei (King, 1970). We staged 234 fixed egg chambers to calculate the frequency distribution of different stages of egg chambers. Further, we also counted the number of egg chambers in each ovariole from these fixed samples (mean number of egg chambers/ovariole (EO) = 5.2±0.09 SEM; n=45).

#### Estimating oogenesis timeline to generate growth curve

We counted the number of ovarioles in a female by dissecting the ovaries from well fed 4-5 days old, mated females in 0.3% PBS-tween. We scored 16 females from two independent replicates (mean number of ovarioles per female(O) =35.63±0.46 SEM, n=16), To obtain average number of eggs laid per female per day, we maintained 4-5 days old females and males in individual pairs in fresh food vials supplemented with fresh yeast paste and recorded the number of eggs laid by individual females in 24 hours. We recorded the fecundity of 8 females from two replicates (mean number of eggs laid per female/day (E)=51.5±2.65 SEM; n=8). This gave us an egg production rate per ovariole (EPR = E/O) (David and Merle, 1968; King, 1970; Spradling, 1993) of 1.45 eggs per day by an individual ovariole (EPR = E/O) From this we estimated the total time required for progression from stage 2 to stage 14 (G = EO/EPR) as 86 hours. Finally, we calculated the growth duration of an egg chamber at a specific stage (Gs) based on the frequency with which it occurred in our samples (eg. since 7.7% of our egg chambers were in stage 8, Gs_8_ = 6.65 h). The frequency of distribution for all stages and their corresponding development time is given in Table S1.

With the estimated duration for the individual stage and the midsection area we fit a standard exponential growth curve: A = 273 e ^0.06t^, where t=time in hours, A=area in µm^2^. From this equation, we calculate the expected time t_0_ (initial age of the egg chamber at the being of the imaging session) for the known area measurements for each time lapse image in Fig. 2G-H and Fig.3F-H.

The increase in egg chamber growth declines past stage 10, we considered the percent of the egg chamber area occupied by the oocyte to fine tune the staging of the egg chambers for yolk uptake and histone transfer experiments (Fig. 4). Therefore, we generated a standard exponential growth curve based on oocyte percent for the stages 8-11 using measured oocyte percent and the estimated duration from fixed samples (n=44). Here is the exponential equation for the oocyte percent: O = 0.318 e ^0.07t^, where t=time in hours, O=oocyte percent. From this equation we calculated the expected time t_0_ (initial age of the egg chamber at the start of the imaging session) from the know oocyte percent of the live imaged samples of Yp1-sfGFP and H2Av-mRFP quantification experiments. We also calculated the correlation for oocyte percent and egg chamber area (Fig. S3A) to make sure egg chamber midsection area and oocyte percent are positively correlated.

#### Yolk proteins quantification

To quantify the sfGFP-tagged yolk protein, we drew the region of interest (ROI) around the oocyte using the H2Av-mCherry signal. Due to our inability to image the full depth of the older egg chambers, we considered one-half of the egg chambers for quantification (top through midsection). We measured the total intensity of GFP for all the z slices across one-half of the egg chamber. Similarly, we measured the total intensities of H2Av-mCherry from the follicle cells to normalize the GFP intensities and obtain normalized total Yp1 intensities over the developmental duration.

#### H2Av-mRFP quantifications

To quantify the total nuclear H2Av-mRFP intensities and volume of the nurse cell nuclei (Fig. 4G-K) we segmented 1-2 posterior nuclei per egg chamber in 3 dimensions using a custom-built FIJI macro which apply Gaussian Blur in 3D (5,5,1) and set an automatic threshold based on Otsu method to generate masks. These masks were further processed by applying erode, dilate, open, close, fill holes functions to refine the segmentation of the nuclear signal across the Z-stacks. Processed masks were converted to ROIs using analyze particles function. These ROIs were used to measure the nuclear area and pixel intensities from unprocessed images.

To estimate how local concentrations change over time (Fig. 4E,G,I-K, S3B) we measured average pixel intensity in 20x20 pixel (428.5 µm) regions in a set of 3 Z-slices (step size 3 µm) starting 9µm below the follicle cell border. Nurse cell cytoplasm was defined as a nuclear free region next to the quantified nucleus within a distance of 100 pixels (17µm) from the oocyte border. “Anterior oocyte cytoplasm” was defined as a nuclear-free region within 100 pixels (17µm) from the border of nurse cells on the oocyte side. “Posterior oocyte cytoplasm” was defined as a region within 100 pixels (17µm) from the posterior border of egg. For quantifying H2Av-mRFP overtime, we considered every fifth time frame. We manually placed the ROI at every time frame considered for analysis to ensure ROI was placed in a relative consistent region across the image duration since the sample moved in all three planes during the imaging intervals. The nurse cell nuclear and cytoplasmic and oocyte cytoplasmic intensities were normalized to follicle cell intensities.

#### 3D surface reconstruction

H2Av-mRFP signal from of the nurse cell nuclei was used to reconstruct the nuclear surfaces, while follicle cell nuclei were used to reconstruct surfaces for the whole egg chambers using the Arivis Vision4D program (version 4.0) (Fig. 2C). An analysis pipeline consisting of blob finder was used to segment nuclear signal based on size thresholding. Further, split threshold was set to avoid merging the of signals or over-segmentation. H2Av-mRFP signal on the surface of the egg chambers was used to train the Arivis machine learning based draw tool. Segmentation errors were corrected manually.

#### Figure preparation

Images were processed in FIJI for figure preparation (Figures 1F,G, 2C,D,F-H, 3F,H, 4A,D,F, S4) using the following functions: Subtract background with a radius of 50-100 pixels, Gaussian blur of radius 0.5-1 pixels, and a Median blur of radius 1-2 (Jackson et al., 2023b). Note, all quantifications were performed on unaltered images. Fig. S4 compares the unprocessed and processed images. Exponential growth curves and graphical presentation of data were prepared out in Python environment. Pictorial presentations and figure panels were prepared in Adobe Illustrator (Version 27.9).

### Statistics

P-values for fecundity and different liquid food diet (Fig. 2B, S1) were derived from a student t-test using Microsoft Excel (Version 16.83). The number of biological replicates are indicated in the figure legends and text.

## ACKNOWLEDGMENTS

We thank Lucy O’Brien and Leslie Koyama for sharing the original Bellymount technology. We thank members of the O’Brien lab, Rocky Diegmiller, members of the Amodeo lab and participants of the GSA DGRC-2023 meeting for constructive comments. We also thank Pat Robison, Ann Lavanway, Bing He, The Dartmouth bio-MT Molecular Interactions & Imaging Core and the Life Sciences Light Microscopy Facility for help with image acquisition and analysis and Robert Robertson for help with the Arduino anesthesia system. We appreciate Yuki Shindo for sharing a *Drosophila* stock (w; Sp/CyO; EndogenousH2Av-mCherry). This work was supported by NIH NIGMS MIRA (1R35GM150853-01) and NIH NIGMS COBRE award (P20-GM113132).

## Author Contributions

Conceptualization: S.B., A.A.; Methodology: S.B.; Validation: S.B., A.A.; Formal analysis: S.B.; Investigation: S.B., AA.; Resources: AA.; Data curation: S.B.; Writing-original draft: S.B., A.A.; Writing-review editing: S.B., A.A.; Supervision: A.A.; Project administration: A.A.; Funding acquisition: A.A.

## Data availability

All relevant data can be found within the article and its supplementary information. The codes for CO_2_ pulsing and imaging analysis will be provided on request.

## MOVIE LEGENDS

<u>Movie 1</u> Fly restraint in Bellymount-PT mount kept on the bench for about 16 hours can lay eggs and stay active.

<u>Movie 2</u> Zoomed-out view of a mount kept at room temperature showing multiple females kept on bench for about 16 hours.

<u>Movie 3</u> A fly that survived 16 hours of imaging is still active and eggs deposited during the imaging period are visible in the mount.

<u>Movie 4</u> View through z stacks across stage 6 egg chamber imaged on the multiphoton microscope for a single time frame (Fig. 2C).

<u>Movie 5</u> 3D reconstruction of two early egg chambers from Fig. 2C, 3D surface with H2Av-mRFP signal

<u>Movie 6</u> 3D reconstruction of two early egg chambers from Fig. 2C, 3D surface without H2Av-mRFP signal

<u>Movie 7</u> Tracking multiple egg chambers for 14 h through Bellymount-PT(Fig. 2G).

<u>Movie 8</u> Tracking a stage 10 egg chamber through nurse cell dumping and dorsal appendage formation during (Fig. 2H).

<u>Movie 9</u> Tracking a young pre-vitellogeneic egg chamber (Fig. 3F).

<u>Movie 10</u> Tracking egg chamber during active vitellogeneic stage (Fig. 3H).

<u>Movie 11</u> Proof of principle movie to show that we can visualize border cell, however speed of the border cell migration is much slower.

<u>Movie 12</u> The egg chamber rotation in mid-stage egg chambers could be visualized in the mid-staged egg chambers along the ovarioles.

<u>Movie 13</u> Older egg chambers also retain their rotation behavior, while egg chambers during nurse cell elimination past stage11 show a distinct flipping behavior rather than a complete rotation.

<u>Movie 14</u> Yp1-sfGFP uptake by an stage 8 egg chamber (Fig. 4A).

<u>Movie 15</u> Early stages of nurse cell dumping and transfer of H2Av-mRFP (Fig. 4D). <u>Movie 16</u> Late-stage nurse cell dumping and transfer of H2Av-mRFP (Fig. 4F)

**Fig.S1.**
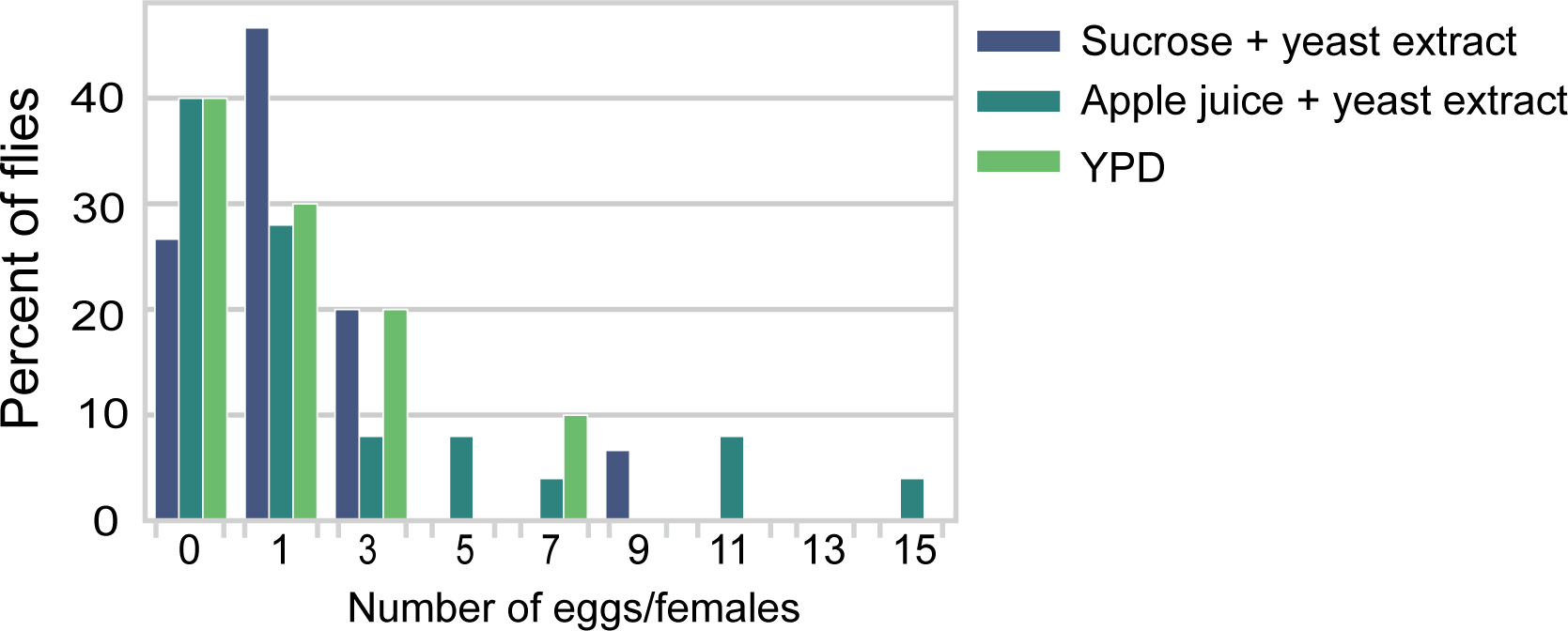
Different liquid foods were tested for their ability to support oogenesis during long term imaging We tested combination of glucose and protein diet in a liquid media to ensure the flies are hydrated, well fed. We used 10% sucrose and yeast extract (n=15), apple juice extract with yeast extract (n=25) and yeast extract peptone dextrose media (YPD) (n=10). We added a few grains of active yeast into each solution. Though we did not observe any significant difference in the average eggs laid by females fed with these diets (p<0.05) maximal egg production occurred on apple juice with yeast extract.

**Fig.S2.**
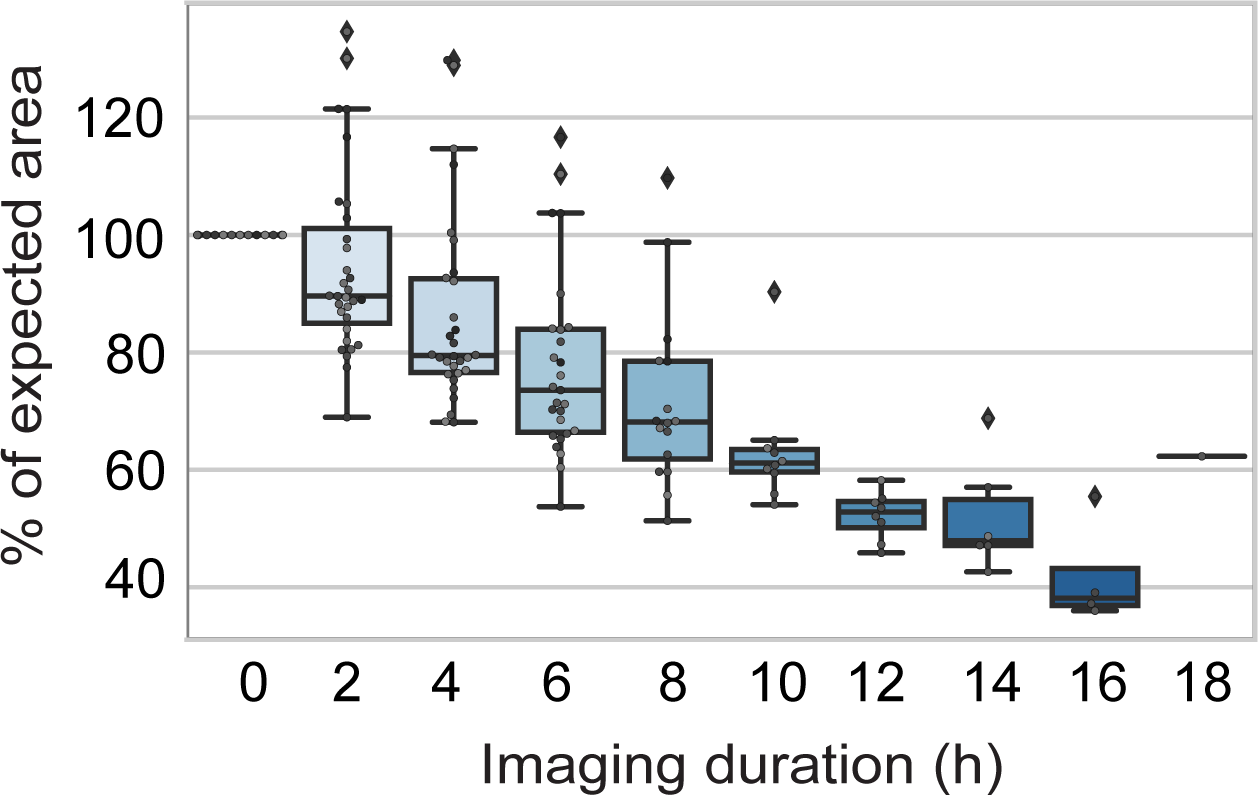
Box plots of the percent of expected area each egg chamber had achieved by the given hour of imaging. Egg chamber growth is slowed in the imaged females compared to the expected values derived from fixed samples (Figure 3B) (n=31, egg chambers stages 4-8).

**Fig.S3.**
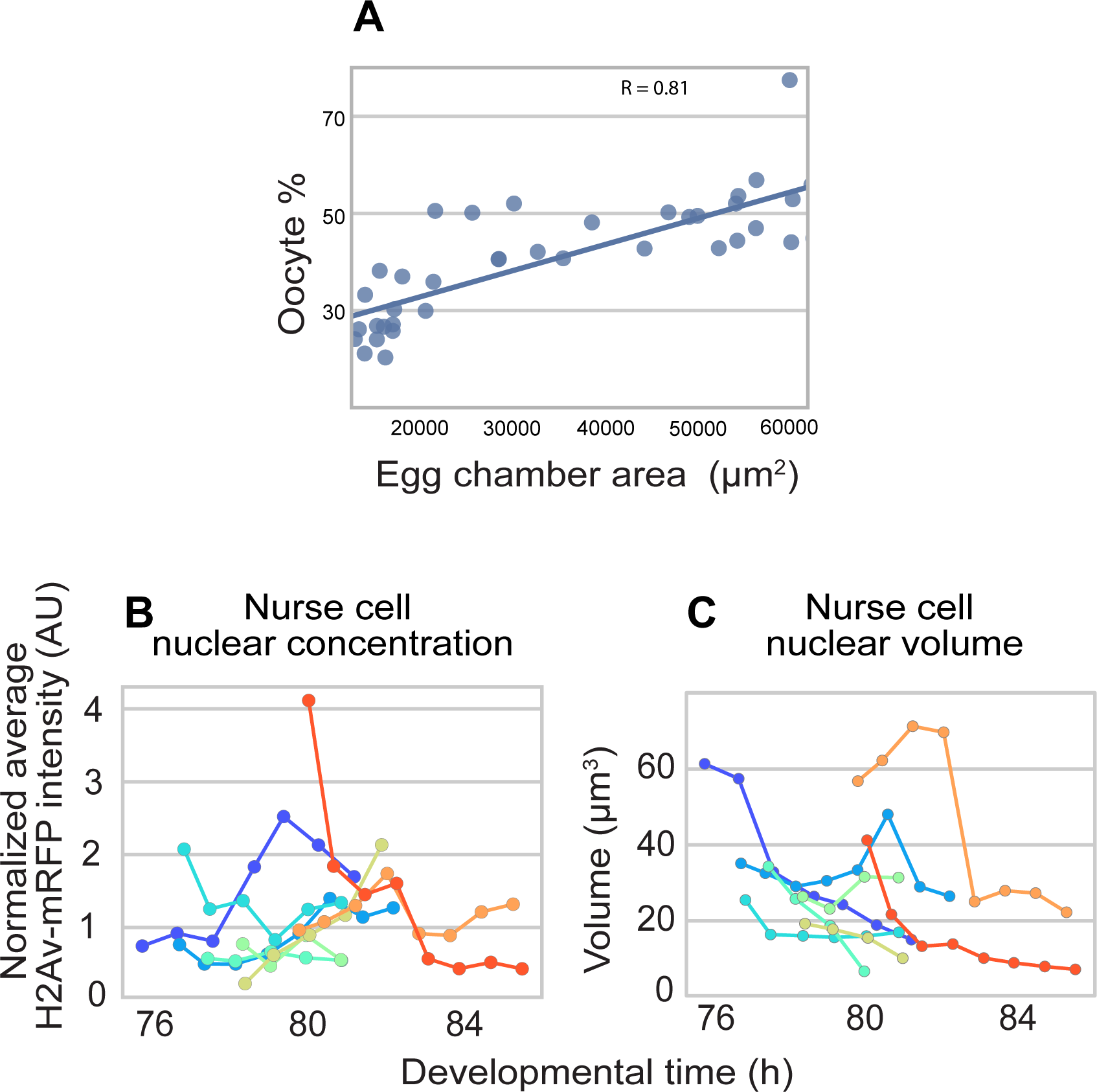
(A) Correlation plot of the oocyte percent against the overall egg chamber area (n=44). We used whole egg chamber size to measure the growth rates (Figure 3). However, we relied on oocyte percent to stage the events that occur during vitellogenesis and dumping (Figure 4). (B and C) Are additional quantifications of the egg chambers in Fig.4 D-K. (B) Normalized average pixel intensity (concentration) of H2Av-mRFP remains relatively constant, or slightly increasing during early nurse cell dumping. (C) Conversely, the total nurse cell nuclear volume falls steadily throughout dumping. This results in the steady decrease in total H2Av-mRFP signal observed in Fig. 4H.

**Fig.S4.**
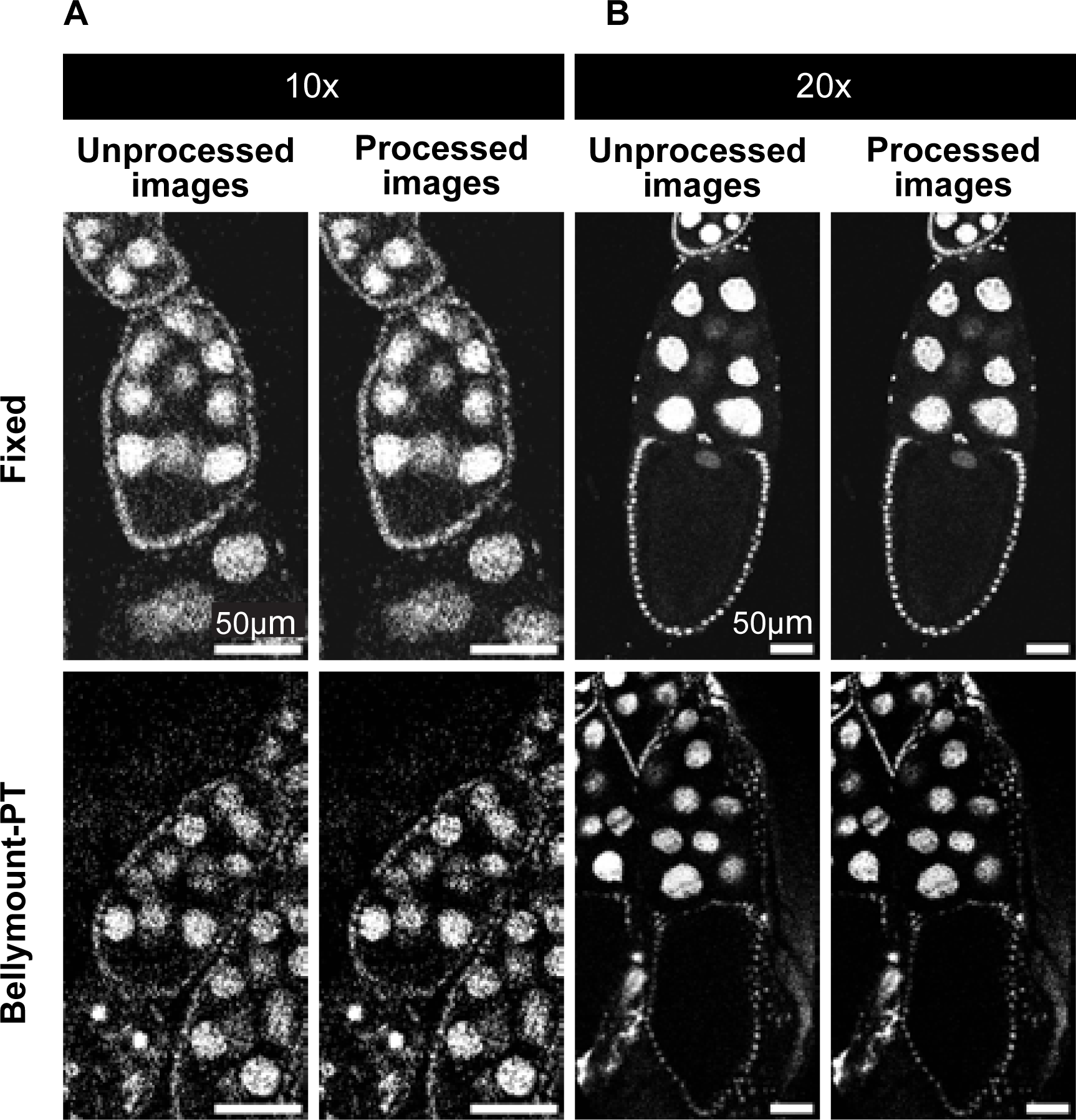
Comparison of unprocessed and processed images for presentation in figures. We used a low degree of filtering to reduce noise. Note that the images used for analysis were not processed. We applied brightness and contrast adjustments, background subtraction, gaussian filter and median filter. (A) is an image captured on 10x objective while (B) from a 20x objective.

**Table S1:** Frequency distribution and developmental duration of individual chambers from stage 2-14, and the total cumulative duration of oogenesis to each stage.

| Stages | Egg chambers size<br>($\mu\text{m}^2$) | Egg chambers | Frequency | Duration /stage (h) | Total cumulative duration of<br>oogenesis' to each stage (h) |
| --- | --- | --- | --- | --- | --- |
| 2 | 621.92±20.94 | 23 | 9.8291 | 8.4854 | 8.4854 |
| 3 | 931.01±26.22 | 29 | 12.3932 | 10.6990 | 19.1845 |
| 4 | 1601.33±56.44 | 31 | 13.2479 | 11.4369 | 30.6214 |
| 5 | 2746.23±82.6 | 23 | 9.8291 | 8.4854 | 39.1068 |
| 6 | 4251.8±145.94 | 31 | 13.2479 | 11.4369 | 50.5437 |
| 7 | 8206.29±283.44 | 26 | 11.1111 | 9.5922 | 60.1359 |
| 8 | 14315.84±694.04 | 18 | 7.6923 | 6.6408 | 66.7767 |
| 9 | 32791.42±2871.94 | 16 | 6.8376 | 5.9029 | 72.6796 |
| 10 | 59036.84±2081.46 | 11 | 4.7009 | 4.0583 | 76.7379 |
| 11 | 63994.96±929.76 | 3 | 1.2821 | 1.1068 | 77.8447 |
| 12 | 72549.9±1917.56 | 9 | 3.8462 | 3.3204 | 81.1650 |
| 13 | 79614.48±920.01 | 2 | 0.8547 | 0.7379 | 81.9029 |
| 14 | 79410.55±1951.13 | 12 | 5.1282 | 4.4272 | 86.3301 |

**Supp. Table 2:**
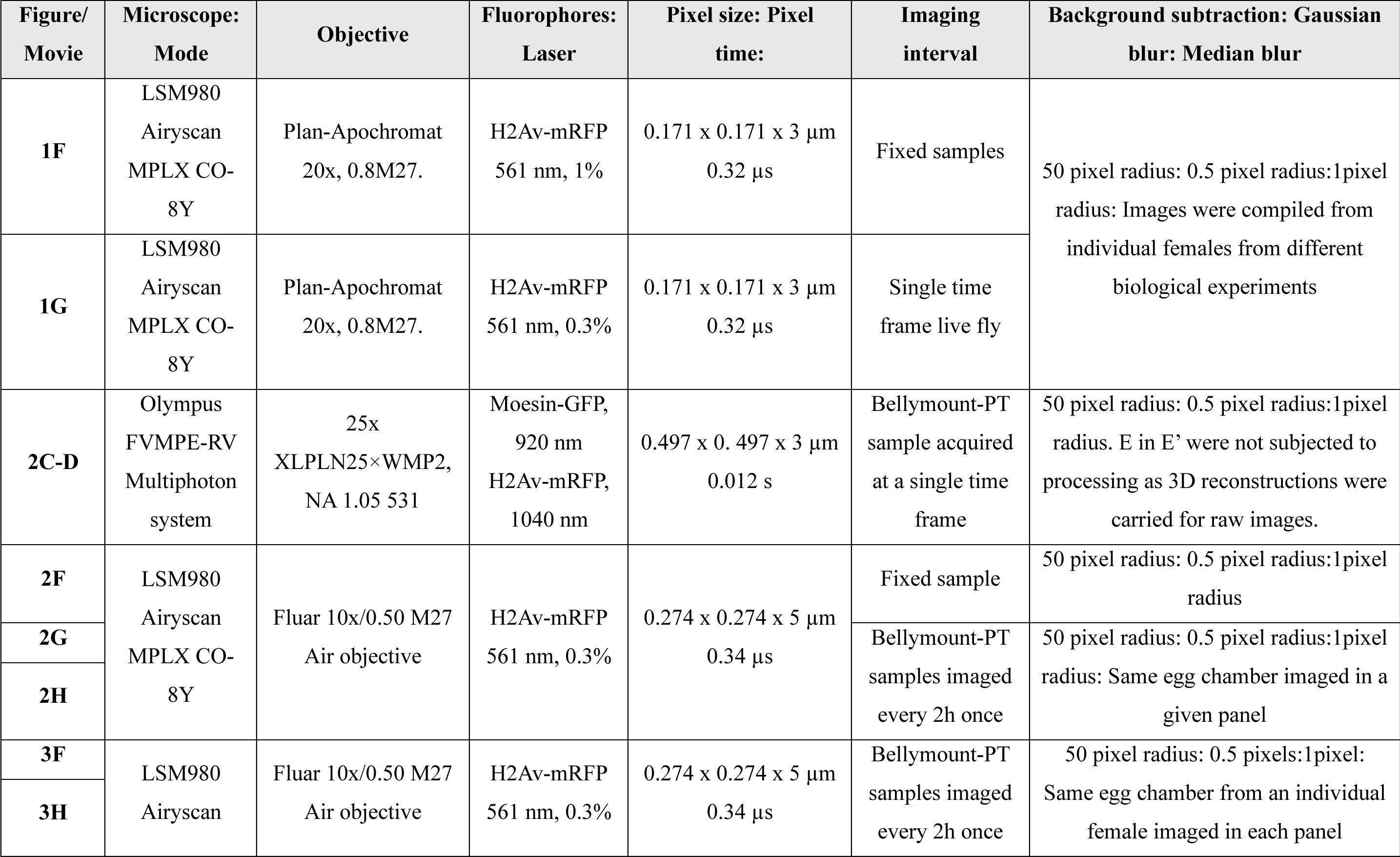
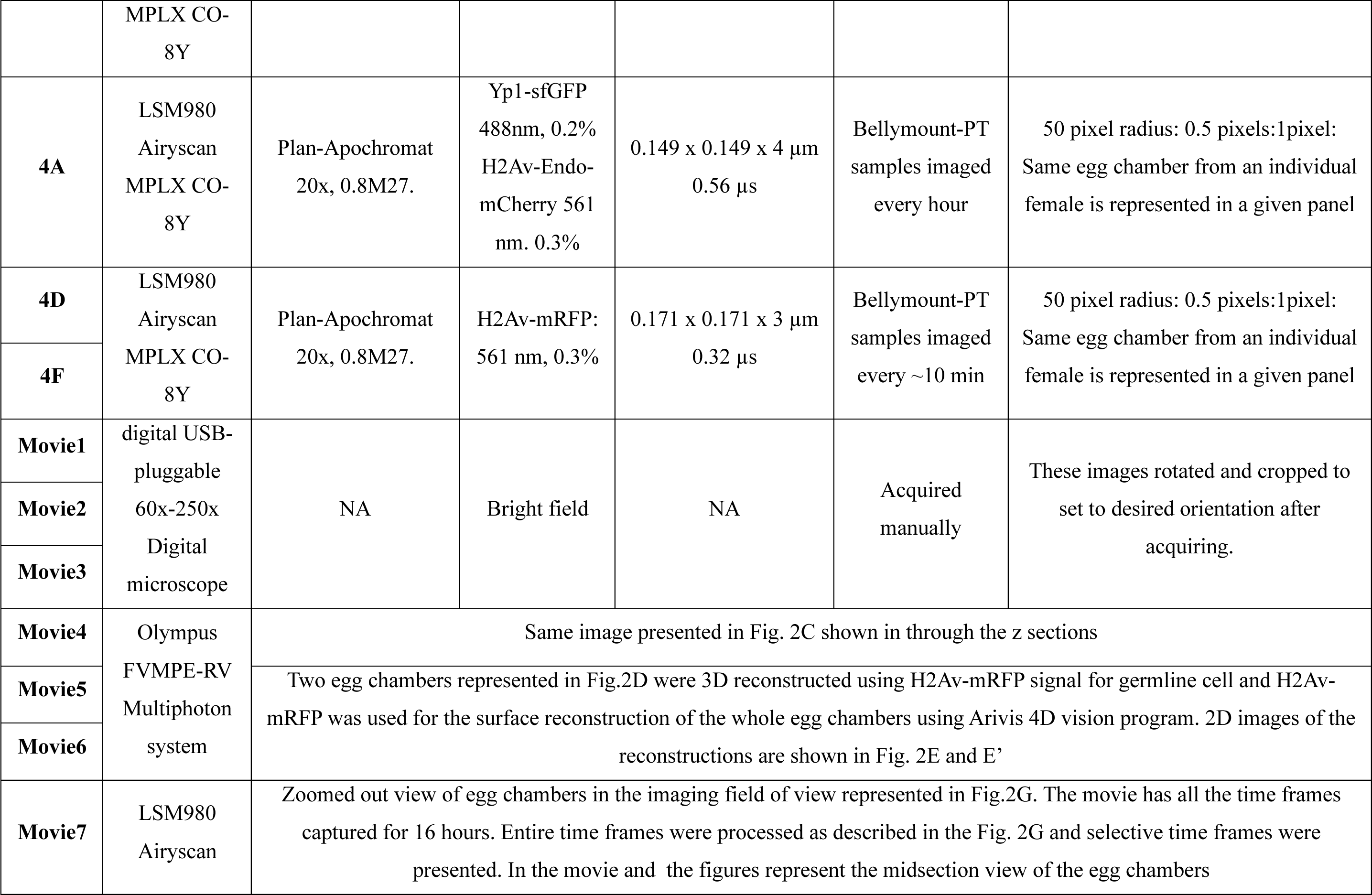
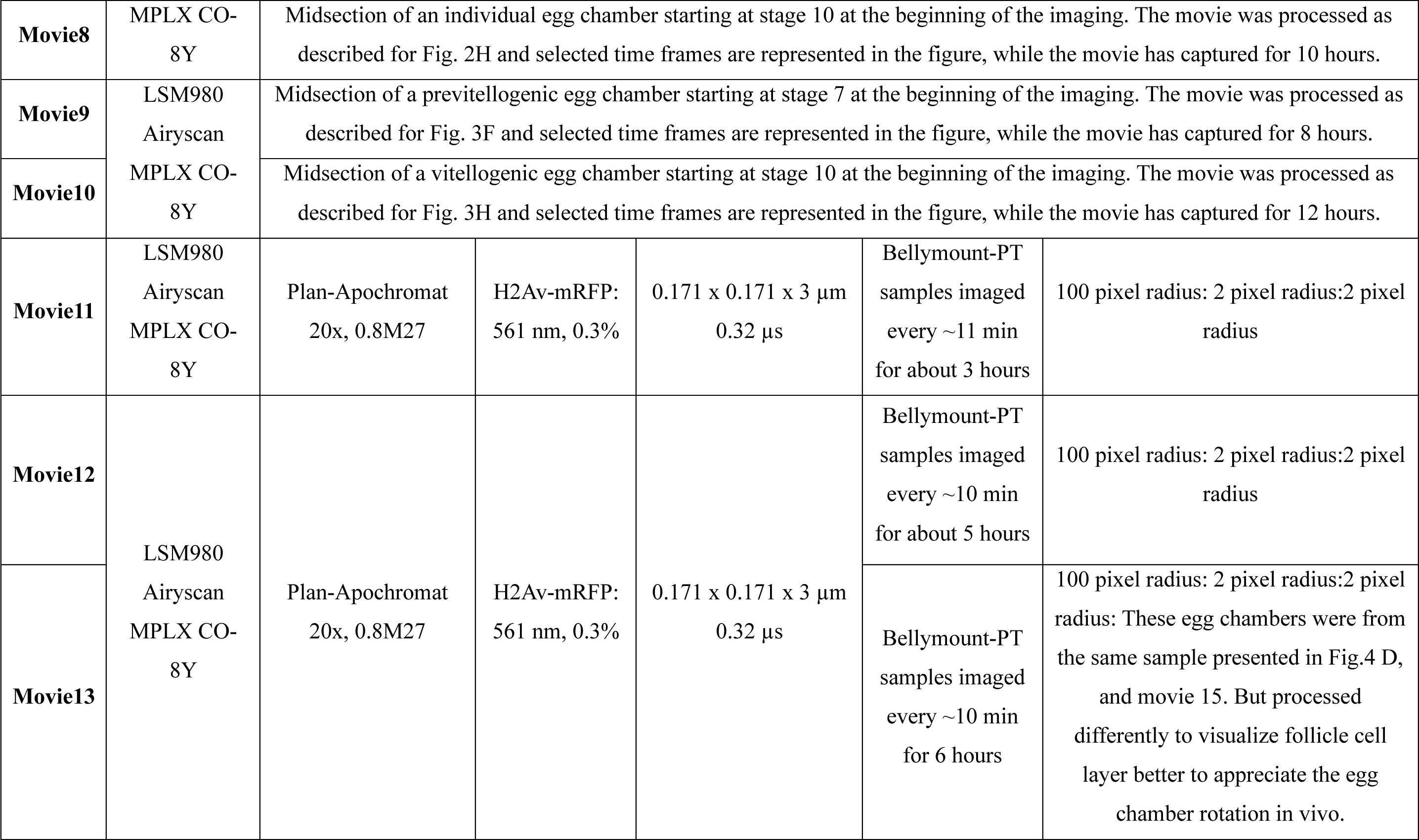
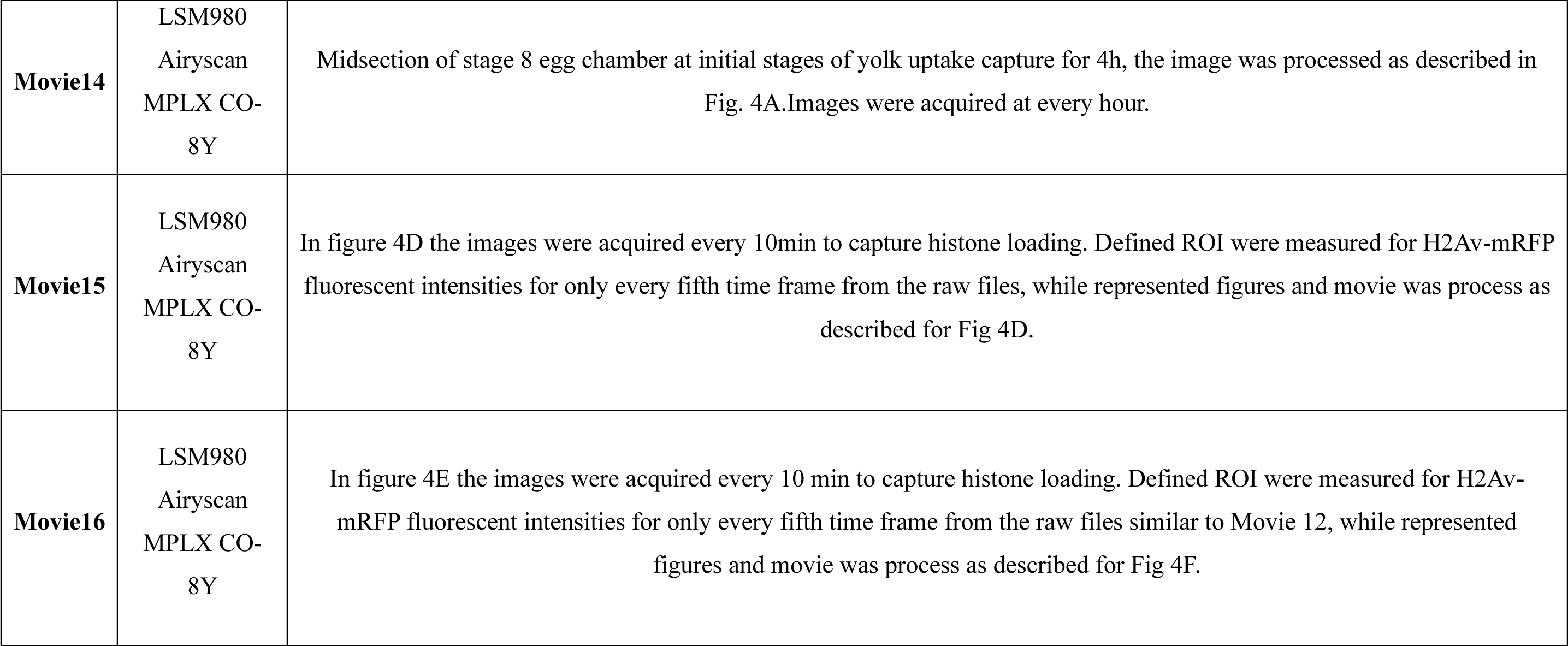
Information related to microscopy and image processing for all images in this paper.

